# Kidney tubular epithelial cell ferroptosis links glomerular injury to tubulointerstitial pathology in lupus nephritis

**DOI:** 10.1101/2022.05.26.493579

**Authors:** Abdel A Alli, Dhruv Desai, Ahmed Elshika, Laurence Morel, Marcus Conrad, Bettina Proneth, William Clapp, Carl Atkinson, Mark Segal, Louis A. Searcy, Nancy D. Denslow, Subhashini Bolisetty, Borna Mehrad, Yogesh Scindia

## Abstract

**Objective:** An appreciation of factors that lead to tubular injury in lupus nephritis is lacking. Iron accumulates in the kidney tubules of nephritic patients and lupus-prone nephritic mice. Ferroptosis is a druggable, iron-dependent form of cell death that has received little attention in lupus nephritis. This study investigated whether intra-renal ferroptosis is a target for intervention in lupus nephritis.

**Methods:** Kidneys of lupus nephritis patients and two spontaneous murine models of lupus nephritis were characterized for ferroptosis using protein, RNA, and lipidomics-based approaches. Susceptibility of heavy chain ferritin (FtH1; an essential iron sequestration protein) deficient proximal tubular epithelial cells (PTECs) was studied using nephrotoxic serum nephritis and FtH1 knockdown human PTECs. The benefit of Liproxstatin-2, a novel second-generation ferroptosis, was evaluated using human PTECs exposed to lupus nephritis patients’ serum.

**Results:** Human and murine nephritic kidneys have the characteristic markers of ferroptosis, such as 4-hydroxynonenal and acyl-CoA synthetase long-chain family member 4, mainly in the tubular segments. Murine kidneys showed impairment in the glutathione synthesis pathway, decreased expression of glutathione peroxidase 4, a glutathione-dependent ferroptosis inhibitor, and characteristic ferroptotic lipid signature. Loss of FtH1 increased PTEC pathology independent of glomerular injury. These findings were recapitulated in human PTECs. Of translational relevance, Liproxstatin-2 demonstrated a prophylactic and therapeutic benefit in mitigating lupus nephritis patient serum-induced PTEC ferroptosis.

**Conclusion:** Our findings highlight tubular cell ferroptosis as a pathological feature in human and murine lupus nephritis and identify ferroptosis inhibitors as potential novel adjunct therapeutics to treat lupus nephritis.

## Introduction

Progressive kidney failure from lupus nephritis occurs in up to 50% of systemic lupus erythematosus (SLE) cases and is associated with increased morbidity and mortality compared with non-lupus nephritis SLE patients(1). Even after immunosuppressive therapies, the rate of complete remission for proliferative disease remains below 50%(2). This is partly due to our limited understanding of the cellular and molecular pathways driving the pathogenesis of lupus nephritis. The vital role of end-organ susceptibility in progression to end-stage kidney disease (ESKD) in lupus nephritis(3, 4) is shifting the current paradigm that views SLE as a disease inflicted by only a disturbed immune system on passive end-organs. Research suggests the existence of non-immune mechanisms of kidney injury in lupus nephritis(5).

The deposition of immune complexes in the glomeruli is thought to initiate lupus nephritis(6). However, the extent of injury to the tubulointerstitial compartment predicts ESKD in lupus nephritis(7). In SLE/lupus nephritis, IgG, anti-dsDNA antibodies, and albumin are over-absorbed by proximal tubular epithelial cells, activating ROS-sensitive pathways that lead to tubular-injury dependent interstitial inflammation(8). Tubulointerstitial inflammation, fibrosis, and atrophy strongly correlate with kidney pathology, independent of the extent of glomerular damage(9) and predict worse outcomes of lupus nephritis(5, 10, 11). Specific delivery of a calcium/calmodulin-dependent protein kinase IV inhibitor to tubular epithelial cells curbed local kidney inflammation in lupus-prone mice without affecting systemic inflammation(12). Collectively, these studies identify tubular epithelial cells as potential target cells to mitigate kidney injury and prevent the progression to ESKD in lupus nephritis. However, an appreciation of factors that lead to tubular epithelial cell injury following early glomerular injury in lupus nephritis is lacking.

Oxidative stress and ROS worsen pathology in human and animal SLE (13) and lupus nephritis,(14). Iron plays a central role in generating ROS(15) and accumulates in the kidneys of both humans(16) and mice with lupus nephritis(17, 18). Multiple receptors on proximal tubular epithelial cells can import transferrin-bound and non-transferrin-bound iron(16). Thus, iron loading in cells can increase their susceptibility to iron-mediated pathology following the loss of glomerular permeability. We, and others, have demonstrated that pharmacological inhibition of kidney iron accumulation mitigates proximal tubular epithelial cell injury and kidney failure in lupus nephritis, even in the presence of pathogenic levels of anti-dsDNA IgG and glomerular immune complexes(17, 19). While these studies establish the role of iron and tubular cells in the pathogenesis of lupus nephritis, the underlying mechanisms by which iron induces this pathology remain poorly defined.

Bioavailable iron reacts with small quantities of hydrogen peroxide generated as part of normal metabolism in the Fenton reaction, producing reactive oxygen species. In addition, iron also serves as a cofactor for lipoxygenases to catalyze polyunsaturated fatty acids (PUFA) peroxidation(20) and enhances lipid peroxides produced by lipid autoxidation(21), the signature of ferroptosis. Ferroptosis is an iron-dependent form of regulated cell death identified by iron-catalyzed damage to the lipid membrane(22, 23). At a mechanistic level, this process involves the iron-dependent accumulation of lipid ROS and depletion of plasma membrane polyunsaturated fatty acids. Growing evidence has identified a central pathophysiological role for ferroptosis in kidney injury(24), nephron loss, and tubular necrosis(25). Lipid peroxidation, the striking feature of ferroptosis, is increased in lupus patients(26). A recent study implicated for the first time ferroptosis in SLE and lupus nephritis(27). This study identified that lupus serum factors like autoantibodies and type I IFNs cooperatively induce ferroptosis in neutrophils but not in lymphocytes and monocytes. Furthermore, selective haploinsufficiency of the ferroptosis inhibitor glutathione peroxidase-4 (GPX4) in mouse neutrophils recapitulated the key clinical features of human SLE. However, whether ferroptosis contributes to kidney tubular pathology (region of iron accumulation) in human and murine lupus nephritis has not yet been investigated, constituting a significant gap in our current understanding of the pathogenesis of this complex disease.

Herein, we observed a clear ferroptosis signature in human lupus nephritis biopsies and kidneys of two spontaneous murine lupus models of different etiologies. Furthermore, we highlight the direct importance of iron sequestration in kidney proximal tubules using the nephrotoxic serum-induced glomerulonephritis and siRNA knockdown studies. Finally, we demonstrate that a novel, second-generation ferroptosis inhibitor Liproxstatin-2, reverses lupus nephritis patient (Class IV) serum-induced ferroptosis in human proximal tubular epithelial cells.

## Materials and Methods

### Mice

All experiments were performed in accordance with the National Institutes of Health and Institutional Animal Care and Use Guidelines and were approved by The Animal Care and Use Committee of the University of Florida. Female MRL/lpr mice were purchased from the Jackson Laboratories, Bar Harbor, USA. (NZW X BXSB) F1 males were bred from parental strains purchased from the Jackson Laboratories. All mice were kept on a standard diet and housed in SPF conditions at the University of Florida. Tissue was collected from 4 and 16 week old (NZW X BXSB) F1 males and 8 and 20 week old female MRL/lpr mice. The transgenic mice deficient in *FtH1* expression only in the kidney’s proximal tubules (*FtH*^*PT−/−*^) were generated as previously described by Bolisetty et.al (28). *FtH*^fl/fl^ males were purchased from Jackson Laboratories. *Pepck*^Cre/wt^ females were gifted by Dr. Volker Haase, Vanderbilt University.

### Induction of nephrotoxic serum glomerulonephritis in mice

Mice were preimmunized intraperitoneally with 200 μg of sheep IgG (Serotec) in Addavax (Invivogen), followed by intravenous injection of sheep nephrotoxic serum (2.5 μL of serum per gram of mouse, Probetex, San Antonia, Tx) 4 d later as described previously(29). Tissue was collected 14 d later.

### Biochemical assays and tissue samples

Proteinuria was assessed by adding 50 μL urine on Siemens Multistix 8 SG dipsticks. Results were graded by color development and converted to absolute numbers as indicated by the manufacturer. Ketamine (120 mg/kg)/xylazine (12 mg/kg) mix was used to anesthetize the mice. Blood was drawn from the axillary vein before euthanasia. Kidney slices were fixed with 10% neutral-buffered formalin for paraffin embedding or with periodate-lysine-paraformaldehyde fixative to be frozen in optimal cutting temperature compound or snap-frozen in liquid nitrogen for subsequent RNA extraction and protein analysis.

### Immunofluorescence

Three-micron cryostat-cut kidney sections were used for the immunofluorescence detection of α smooth muscle actin (αSMA), F4/80+ macrophages, CD4+ T cells and immune complexes. Details of staining and antibodies is provided in the Supplemental Methods.

### Immunohistochemistry

Paraffin-embedded human biopsies or mouse kidney sections (3 μm) were stained for acyl-CoA synthetase long-chain family member 4 (ACSL4) or 4-hydroxynonenal (4-HNE) using standard protocols. Details of immunostaining are described in Supplemental Methods section.

### Cell Culture and siRNA knockdown studies

HK-2 cells, a normal human proximal tubular cell line (ATCC) were maintained in ATCC recommended keratinocyte serum-free medium with the addition of bovine pituitary extract (0.05 mg/mL) and human recombinant epidermal growth factor (5 ng/mL). All experiments were carried out in a medium containing 5% serum from lupus nephritis (Class IV) patients or healthy donors. 2 × 10^5^ cells were pretreated with 100 ng/mL Liproxstain-2 (a proprietary new generation ferroptosis inhibitor by ROSCUE Therapeutics with improved metabolic stability in human microsomes) or DMSO for 12 hrs before serum treatment. In some experiments liproxstain-2 was added after 1 or 4 hours after the addition of patient serum. All data were analyzed 24 hrs after addition of patient or healthy donor serum.

For gene knockdown studies, silencer select siRNA for human FtH1 and negative control siRNA (4392421 and 4390843 respectively, ThermoFisher Scientific) were used. Experimental details are provided in Supplemental Methods section.

### Preparation of Kidney and Cell Extracts, and Western Blotting

Western blotting was performed as described in our previous publication (19). Details of tissue digestion, antibodies and the dilutions are provided in the Supplemental Methods section.

### Detection of serum IgG, IgG2a, and anti-dsDNA antibodies

Mouse serum IgG and IgG2a were detected using commercially available ELISA kits (ThermoFisher) as per manufacturers’ instructions. Anti-dsDNA IgG was measured in 1:100 diluted serum using plates coated with 50 μg/ml dsDNA. Bound IgG was detected using alkaline phosphatase-conjugated anti-mouse IgG (Jackson ImmunoResearch, West Grove, PA) diluted 1:1000. Relative units were standardized using serial dilutions of a positive serum from SLE123 triple congenic mice, setting the 1:100 dilution reactivity to 100 U(30).

### RT-PCR

For RNA isolation, frozen tissues or live cells were resuspended in RLT buffer (Qiagen, Valencia, CA) and homogenized using the TissueLyser system or Qiashredder (Qiagen). RT was performed as described in our previous publication (19), the details of which are provided in the Supplemental Methods section.

### Lipid extraction and liquid chromatograhpy-mass spectrometry

Lipids were extracted with a modified Bligh-Dyer protocol(31). The details of sample preparation and conditions for performing liquid chromatography-mass spectrometric analysis are provided in the Supplemental Methods section.

### Study approval

The study was performed per the Declaration of Helsinki, under protocols approved by the University of Florida Institutional Review Board (IRB#201601162 and IRB#201601170) after written informed consent was received from participants.

### Statistics

Statistical significance was determined by applying a Mann-Whitney test for groups not passing the normality test. 1-way and 2-way analysis of variance (ANOVA) with Tukey’s or Holm-Šídák’s multiple comparisons test were used to compare more than 2 groups of experimental conditions and represented as mean ± SEM. All analyses were performed using GraphPad Prism 9 (GraphPad Inc, San Diego, CA).

## Results

### Tubular injury is a prominent feature in lupus nephritis

Glomerular immune complexes were observed at 8 weeks (**Figure 1A**), whereas proteinuria was negligible (8 weeks,10±6 mg/dL). However, intense glomerular and tubular immune complexes (**Figure 1B, white arrows indicate tubular immune complexes**) and severe proteinuria (20 weeks, 1775±147 mg/dL) were a feature in 20-week-old mice. There was no histological evidence of injury (**Figure 1C, Supplemental Figure 1A)** at 8 weeks. In comparison H&E staining showed a high degree of cellular infiltrates surrounding the injured glomeruli and tubulointerstitial regions (**Figure 1D**). Tubulointerstitial injury was a distinct and prominent feature in diseased kidneys. Multiple atrophic tubules with loss of brush border and enlarged basement membrane, blebbed tubules, and interstitial fibrosis were observed in nephritic kidneys (20-week-old, **Figure 1D**). Further, periodic acid Schiff’s (PAS)-stained kidney sections of nephritic mice revealed classic lupus nephritis features such as severely sclerosed glomeruli with crescents (**Supplemental Figure 1B**). This indicates that presence of immune complexes does not result in observable kidney injury. Sensitive markers of proximal tubular injury, NGAL and KIM-1, were not detected at 8 weeks but were significantly elevated at 20 weeks of age (**Figure 1E-F)**, corroborating the tubulointerstitial injury seen in our histological analysis. Immunofluorescence staining of the kidneys revealed immune cell infiltrates composed of macrophages (**Supplemental Figure 1C-D**), and CD4 T cells were observed in the peritubular regions (**Supplemental Figure 1E-G**). Thus, while glomerular immune complexes are observed when kidney injury is asymptomatic (8-weeks), severe tubular injury is a distinct feature during the nephritic stages (20-weeks) and contributes to overall kidney pathology in lupus nephritis.

**Figure 1.**
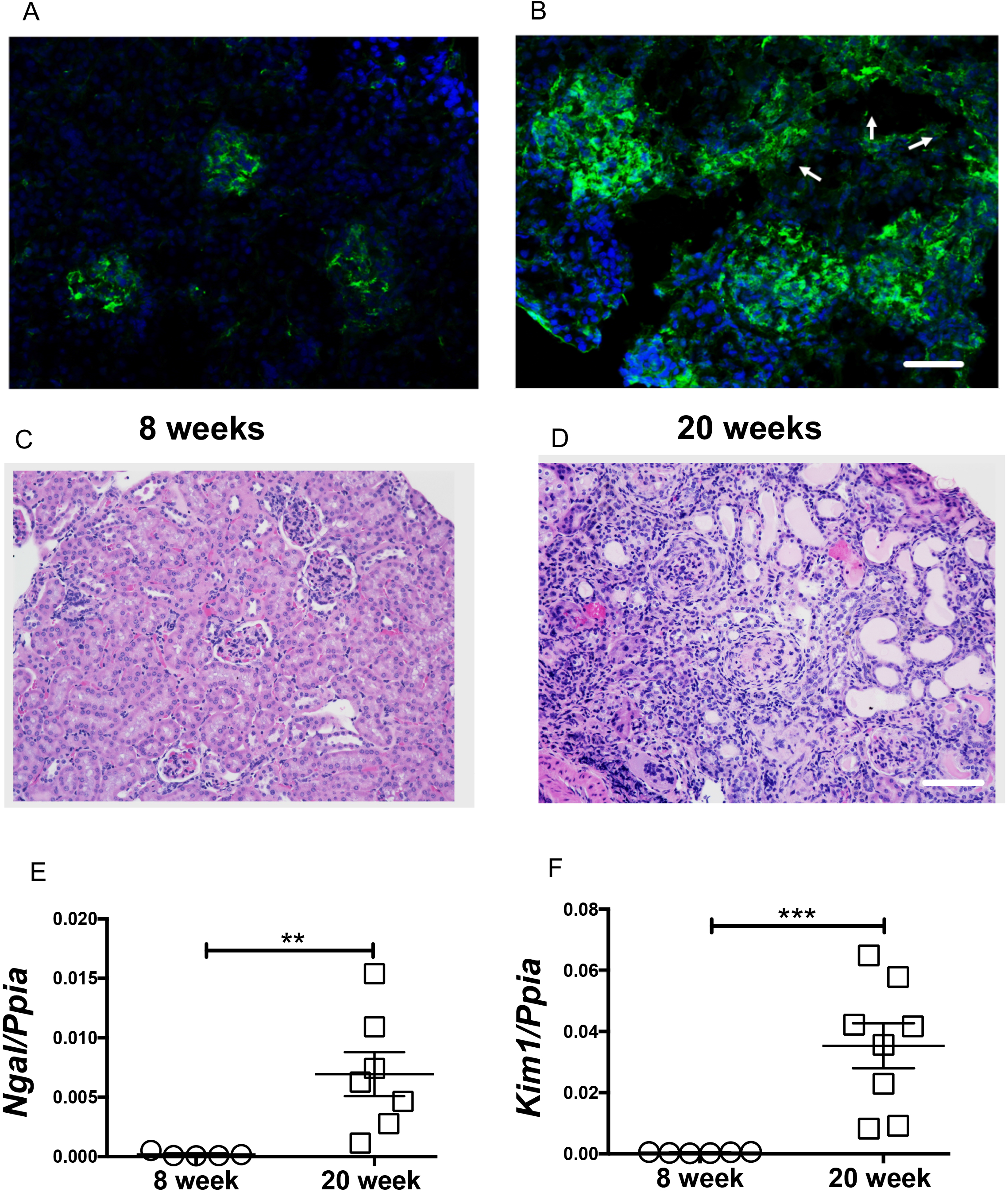
Tubular injury is a distinct feature in lupus nephritis. Glomerular immune complexes were observed in 8-week-old female MRL/lpr mice (A). At 20-weeks of age, glomerular and tubular immune complex deposits are evident (arrow points to the tubules) (B). Compared to 8-week-old female (C), the renal histology (H&E) at 20 weeks showed severe periglomerular and interstitial immune infiltrates. Along with the traditional glomerular injury, large number of a-nuclear tubules, tubular cast are also evident (D). Scale bar = 50 μm and 100 μm. The tubular injury marker Ngal and Kim-1 were significantly elevated in 20-week-old nephritic mice (E-F). Statistical significance was determined by 2-tailed Mann-Whitney test and represented as mean ± SEM. **P < 0.01, ***P < 0.001.

### Iron metabolism dysregulation in the kidneys of nephritic mice

We compared kidney iron content, changes in expression of genes and proteins associated with iron metabolism in non-nephritic and nephritic female MRL/lpr mice. No iron was detectable by Perl’s staining in non-nephritic mice (**Figure 2A)**. However, substantial iron deposits were observed in the tubular cells of nephritic mice (**Figure 2B**). Though severely injured, glomeruli were devoid of iron deposits. Kidney gene expression for *Zip8* and *Zip14*, which encode for SLC39A8 and SLC39A14, the metal transporters that mediate non-transferrin-bound iron uptake into proximal renal tubular cells(32), were significantly decreased in nephritic mice (**Figure 2C-D**). There was a concomitant increase in protein expression of ferritin heavy chain (FtH1) (**Figure 2E-F**). FtH1 sequesters iron for storage within its core(33, 34). Iron deposits in the interstitium and tubular compartment, lower expression of kidney tubular iron importers, and increased expression of FtH indicate iron accumulation in nephritic kidneys selectively in the tubules and interstitium.

**Figure 2.**
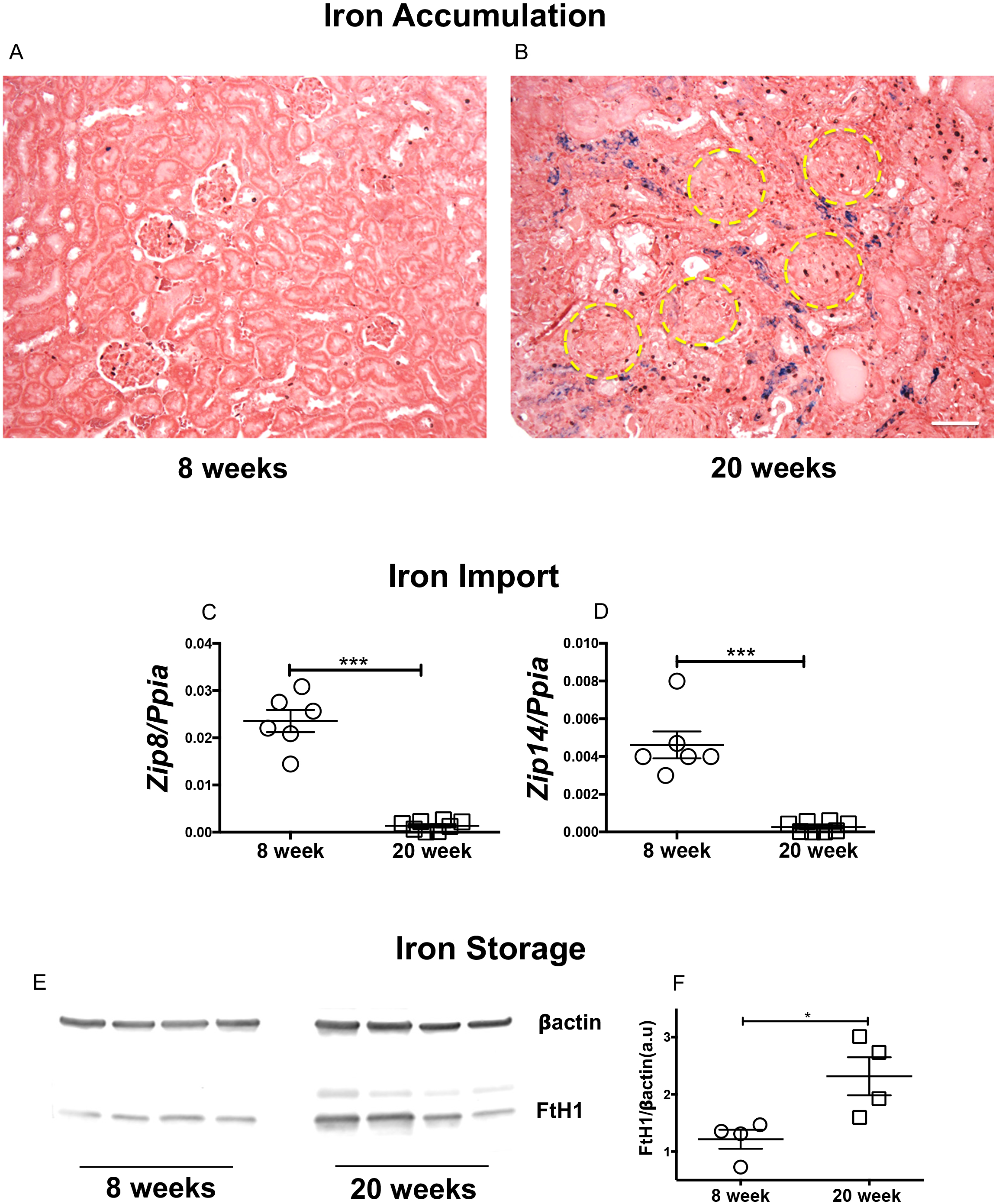
Iron accumulates in the renal tubular segments of nephritic mice and accompanied by significant changes in iron import and storage machinery. Formalin fixed kidney sections from 8- and 20-week-old female MRL/lpr mice were stained for Perls detectable iron, most of which was detected in the interstitium and tubular segments of nephritic mice (A-B). Scale bar = 50 μm. Though severely injured, the glomeruli (yellow circles) were devoid of iron deposits (A-B). Renal gene signature of iron importers, Zip8 and Zip14 was reduced in 20-week-old nephritic female MRL/lpr mice (C-D). Increased renal iron was associated with an increase in expression of heavy chain ferritin (FtH), the endogenous iron sequestration protein (E-F). Statistical significance was determined by 2-tailed Mann-Whitney test and plotted as mean ± SEM. *P < 0.01, ***P < 0.0001.

### Lupus nephritis kidneys display oxidative stress and a ferroptosis gene signature

Ferroptosis is an iron-dependent form of oxidative injury primarily studied in acute tubular injury in the kidneys (24, 35, 36). While iron deposits are documented in lupus nephritis, the occurrence of ferroptosis in murine kidneys in the setting of lupus nephritis has not, to our knowledge, been reported. Compared to young non-nephritic MRL/lpr female mice, we observed an upregulation of NAD(P)H Quinone Dehydrogenase 1 (*Nqo1*), heme oxygenase 1 (*Hmox1*), and downregulation of thioredoxin 1 (*T*x*nrd1*), genes associated with protection against oxidative stress in the kidneys of females (**Supplemental Figure 2A-C**). In addition, increased oxidative stress was associated with a ferroptosis gene signature (**Supplemental Figure 2D-F**).

### Lupus nephritis is associated with impairments in kidney glutathione synthesis and increased expression of ferroptosis core proteins

Glutamate cysteine ligase (GLC) catalyzes the ligation of glutamate and cysteine. As such, is the rate-limiting enzyme in glutathione synthesis(37). GLC comprises two independent gene products, the glutamate-cysteine ligase-catalytic (*Gclc*) and glutamate-cysteine ligase-modifier (*Gclm*) subunits. We found lower gene expression of both *Gclc* and *Gclm* in nephritic MRL/lpr female mice (**Figure 3A-B**). GPX4, a glutathione-dependent ferroptosis inhibitor(38), and ACSL4, a ferroptosis promoter, are two core proteins that may determine the cell’s sensitivity towards ferroptosis(39). However, their expression in lupus nephritis is unknown. The protein levels of GPX4 were lower in the nephritic kidneys (**Figure 3C-E**), coinciding with a higher expression of ACSL4 (**Figure 3D-F**). Furthermore, nephritic kidneys had a higher immunoreactivity to 4-HNE, an established lipid peroxidation marker (**Figure 3G-H**). 4-HNE was primarily observed in the kidney tubules, a region of iron accumulation (**Figure 2B)**.

**Figure 3.**
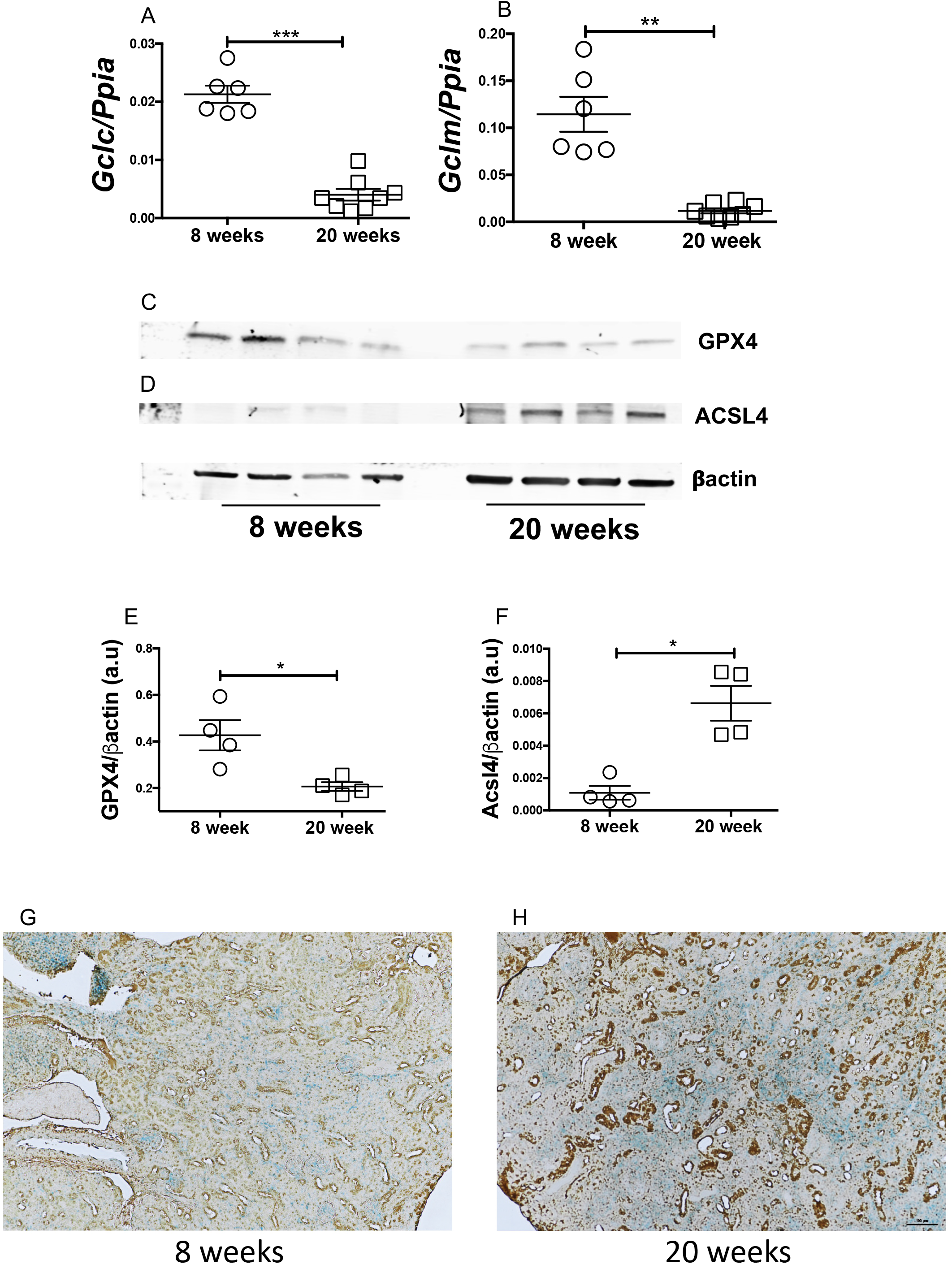
Lupus nephritis is associated with decreased expression of genes involved in glutathione biosynthesis pathway and an increased ferroptosis protein signature. Nephritic female MRL/lpr mice show lower expression of Gclc (A) and Gclm (B), gene products that constitute glutamate cysteine ligase (GCL), the activity of which determines de novo glutathione synthesis. Compared to 8-week-old MRL/lpr females, 20-week-old nephritic females had significantly lower expression of GPX4, the glutathione dependent ferroptosis inhibitor (C and E), whereas the expression of ACSL4, the ferroptosis executioner was significantly elevated (D and F). Nephritic kidneys of MRL/lpr females stain more intensely for 4-HNE, a lipid peroxidation marker (G, H). Most of the staining is in the tubular segments, the area of iron accumulation. Representative images are shown. Scale bar = 50 μm. Statistical significance was determined by 2-tailed Mann-Whitney test. Data are plotted as mean ± SEM. *P < 0.05, **P < 0.001, ***P < 0.0001.

### Lipidomics identifies a distinct profile and pattern in nephritic kidneys

Using a semi-quantitative LC-MS/MS method, we detected several lipid classes and species in non-nephritic and nephritic kidneys. **Figure 4A** is the representative normal phase LC-MS chromatogram for six major classes of phospholipids found in the kidneys. The phosphotidylethanolamine (PE), PE(P-18:0/20:4), PE(P-18:1/20:4) and PE(P-18:0/20:4), which are the major storage depots for arachidonic acid were significantly increased in nephritic mice (**Figure 4B-D**). We also found a significant increase in the esterification of the sn-2 chain of PE with adrenic acid (C22:4) (P-18:0/22:4), the preferred substrate for lipid peroxidation in the nephritic kidneys (**Figure 4E)**. This agrees well with the increased expression of ACSL4 and 4-HNE staining in nephritic kidneys. We also observed a differential distribution of lysophosphatidylethanolamines (LPE), phosphatidylethanolamines (PE), alkyl-phosphatidylethanolamine PE(O), and alkenyl-phosphatidylethanolamine PE(P) between non-nephritic and nephritic kidneys as depicted in the heat maps (**Supplemental Figure 3A-D**). The heat maps in supplemental figure 2A-D are represented as a means of pattern/profile recognition, utilizing a color-code display of most differentially expressed within lipids of each class in Supplemental figure 3E.

**Figure 4.**
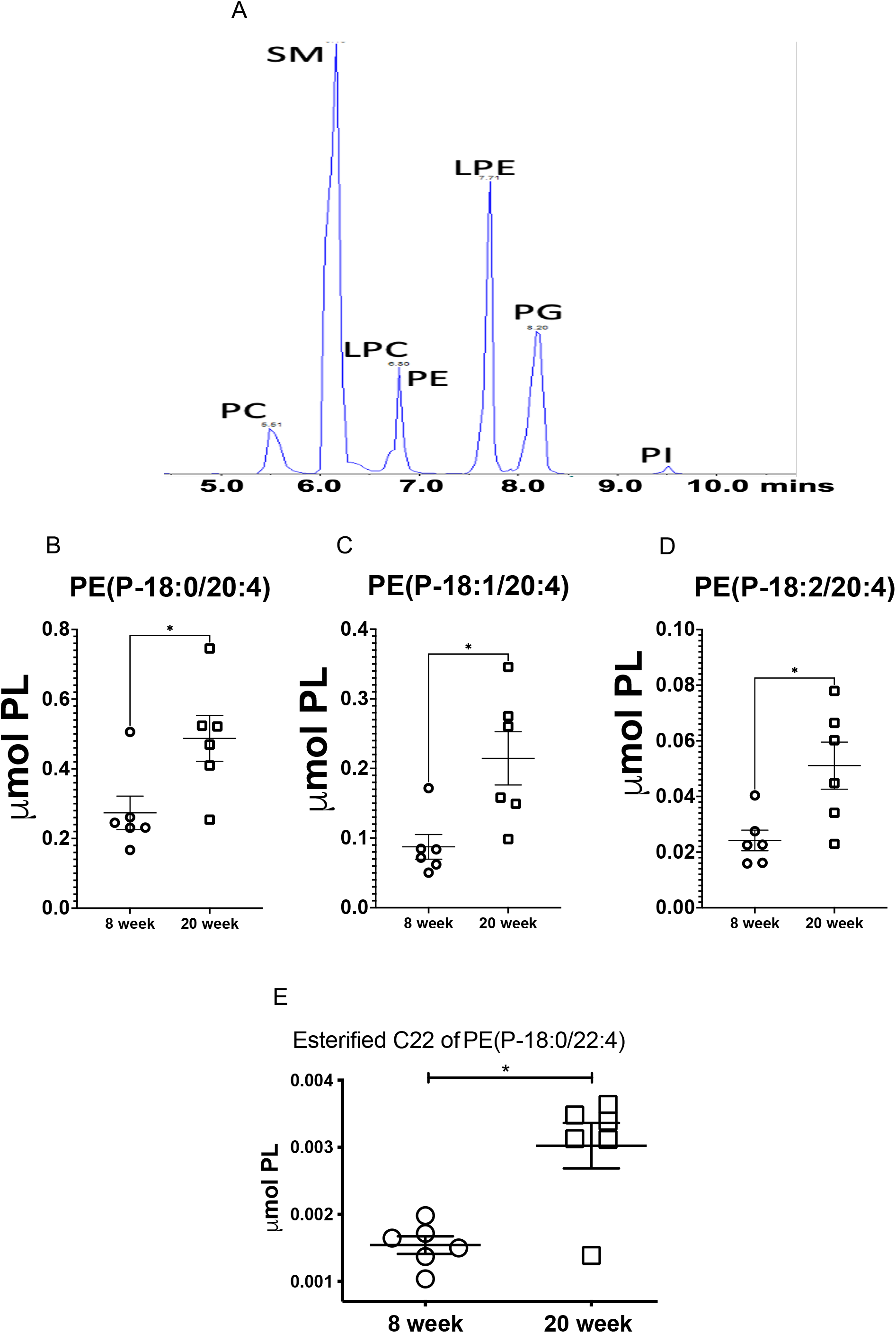
Differential renal lipid profile and increased esterification of phosphatidylethanolamine (PE) SN1 chain in nephritic mice. 8-week-old (non-nephritic) and 20-week-old (nephritic) MRL/lpr female kidneys were analyzed by semi-targeted LS-MS for their lipid profile and content. Representative normal phase LC-MS/MS chromatogram for six major classes of phospholipids: phosphatidylcholine (PC), sphingomyelin (SM),lysophophatidylcholine(PLC),phosphotidylethanolamine(PE),lysophosphotidylethanolamin e (LPE), phosphotidylglycerol (PG), and phosphotidylinostitol (PI) (A). Nephritic mice (20-week-old) had significantly higher concentrations of PE(P-18:0/20:4) (B), PE(P-18:1/20:4) (C) and PE(P-18:0/20:4) (D), the major storage depots for arachidonic acid. Nephritis was also associated with an increase in esterified C22 chain of PE (P-18:0/22:4), the preferred substrate for ferroptosis (E). (n = 6 each). Statistical significance was determined by 2-tailed Mann-Whitney test. Data are plotted as mean ± SEM *P < 0.05.

### Kidney tubular injury and ferroptosis feature in lupus nephritis of different etiologies

Male (NZW X BXSB) F1 mice carry two copies of the *Tlr7* gene and develop proliferative nephritis by 16 weeks age with clinical features of lupus (**Supplemental Figure 4A**)(40). While not as severe as MRL/lpr females, the males presented renal pathology with glomerular and tubular injury. Increased gene expression of *Ngal* and *Kim1* (**Supplemental Figure 4B-F**) indicated damage to the proximal tubular cells with elevated oxidative stress (**Supplemental Figure 4G-I**). These mice also had reduced gene expression of both *Gclc* and *Gclm*, indicating an impaired glutathione synthesis pathway with concomitant decrease in the protein expression of GPX4 (**Supplemental Figure 5A-D**). In addition, we also observed an increased gene expression of *Acsl4* and *Aimf2*, which indicated a ferroptosis signature (**Supplemental Figure 5E-F**). These observations collectively identify tubular injury, impaired glutathione synthesis, and ferroptosis as a common denominator in models with different lupus nephritis etiology.

**Figure 5.**
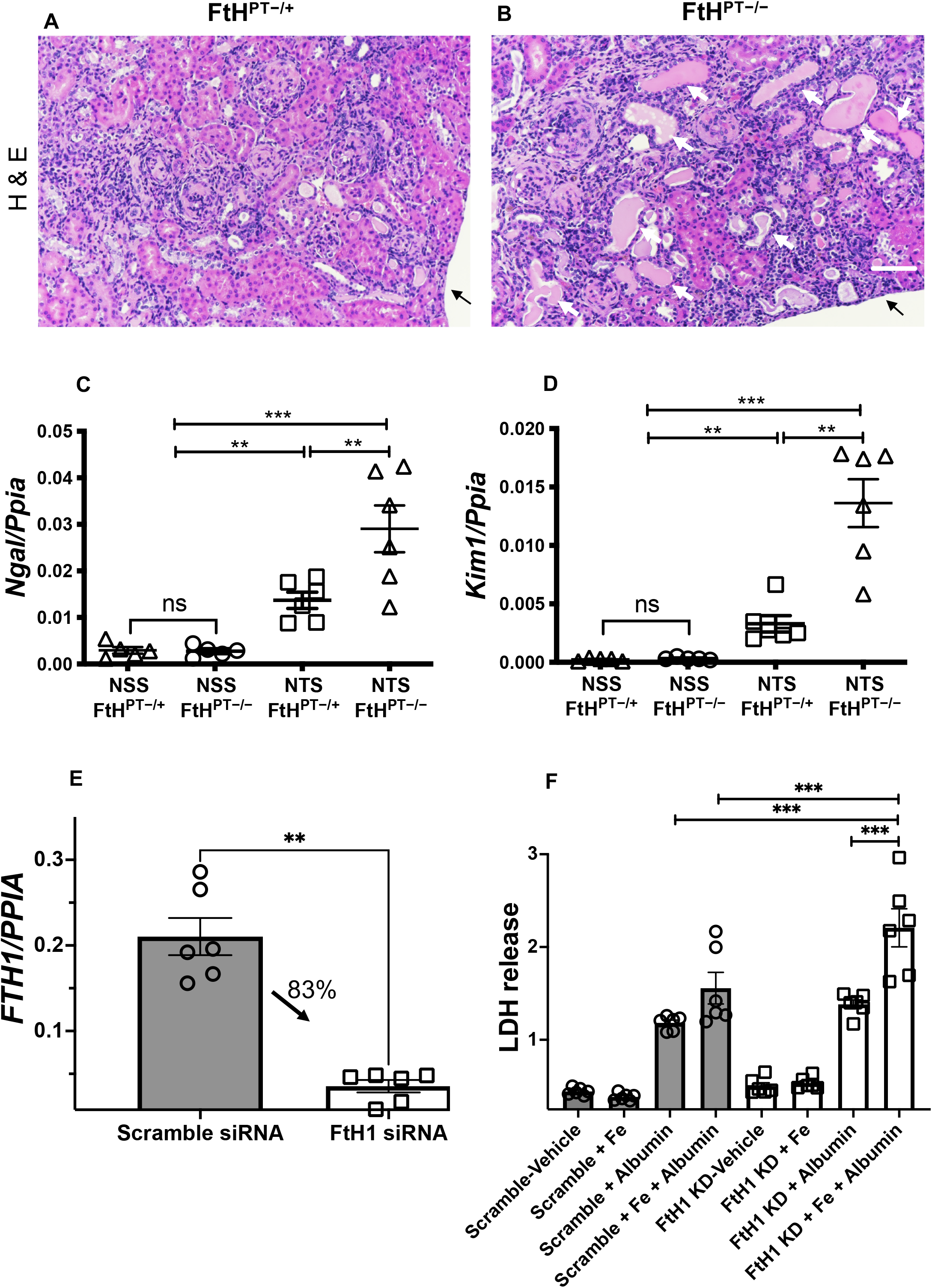
Loss of FtH1 in proximal renal tubules results in increased tubular injury following glomerulonephritis. 12-week-old female FtHPT-/- or FtHPT-/+ mice, were pre-sensitized with CFA (100ug), and four days later were injected i.v., with 100 uL normal sheep serum (NSS) or nephrotoxic sheep serum (NTS). Kidneys were analyzed 14 days later. Hematoxylin and Eosin (H&E) staining revealed inflammatory infiltrates in both the groups. However, compared to nephrotoxic serum injected FtHPT-/+ mice (litter mate controls), FtHPT-/-mice had more tubular epithelial cell necrosis (dark pink fragmented cytoplasm with no nuclei) and denudation of the basement membrane (white arrows). Tubular casts, dilatation, luminal debris were also obvious in the FtHPT-/-(A-B) Arrow denotes end of the section. Scale bar 100 uM. The observed renal pathology was supported by increased expression of proximal tubular injury markers Ngal (C) and Kim1 (D) in FtHPT-/-mice. 2X105 HK-2 cells treated with scramble siRNA or siRNA to H-ferritin and this resulted in ∼ 80% knockdown of H-ferritin gene (E). Following knockdown (83% after 52 hrs), cells were treated with vehicle, Ferrous sulphate (200 μg/mL), or human albumin (20 mg/mL) and the supernatants were analyzed for LDH levels after 12 hrs. Vehicle and Fe treated medium had comparable level of LDH in scramble or FtH1 knock down (FtH1 KD) groups. Albumin significantly but comparable induced cell death in both groups. However, concomitant treatment with Fe and albumin significantly increased in cell death in FtH1 KD HK-2 cells (F). Data was analyzed using 1-way and 2-way ANOVA with Holm-Šídák’s multiple comparisons test and represented as mean ± SEM. **P < 0.001, ***P < 0.0001.

### Impaired iron handling in kidney proximal tubular cells worsens tubular injury in the setting of primary glomerulonephritis

The above observations highlight the importance of iron and ferroptosis in spontaneous lupus nephritis. However, whether disrupted tubular iron metabolism can directly affect outcomes of primary glomerulonephritis is unknown. Proximal tubular epithelial cells have a high abundance of ferritin heavy chain 1 (FtH1), a key iron sequestering protein(41). To elucidate the importance of FtH1 expression in proximal tubular epithelial cells and thus iron metabolism in glomerulonephritis, we induced nephrotoxic serum-induced glomerulonephritis in transgenic mice deficient in *FtH1* expression only in the proximal tubules of the kidney (*FtH*^*PT−/−*^). The genotype of the *FtH*^*PT−/−*^ and *FtH*^*PT−/+*^ mice is shown in **Supplemental Figure 6A**. Fourteen days following induction of glomerulonephritis, there was comparable proteinuria and glomerular immune complex deposits in *FtH*^*PT−/−*^ and *FtH*^*PT−/+*^littermate controls (**Supplemental Figure 6B-D**). In this induced model, as in spontaneous lupus nephritis, the glomeruli were devoid of observable iron deposits (**Supplemental Figure 6E-F**). However, following the injection of nephrotoxic serum, dense but comparable Perl’s detectable iron deposits were observed in both the *FtH*^*PT−/−*^ and *FtH*^*PT/+*^ mice (**Supplemental Figure 6E-F, inset**). We did not observe any detectable iron deposits in normal sheep serum injected *FtH*^*PT−/−*^ and *FtH*^*PT−/+*^ mice. Hematoxylin-Eosin staining revealed extensive immune infiltrates in *FtH*^*PT−/−*^ and *FtH*^*PT−/+*^ mice (**Figure 5A-B)**. However, compared to nephrotoxic serum injected *FtH*^*PT-/+*^ littermate controls, *FtH*^*PT−/−*^mice had significantly more tubular epithelial cell necrosis (dark pink fragmented cytoplasm with no nuclei) with denudation of the basement membrane, tubular casts, dilatation, luminal debris (**Figure 5A-B**). Increased gene expression of proximal tubular injury markers *Ngal* and *Kim1* in the nephrotoxic serum injected *FtH*^*PT−/−*^ mice supported the histopathology (**Figure 5C-D)**. To further validate the importance of proximal tubular epithelial cell FtH1 in increased iron and albumin (the two conditions observed in glomerulonephritis), we knocked down the FtH1 gene by 83% in HK-2 cells (human proximal tubular epithelial cells) (**Figure 5E**). Knocking down FtH1 in HK-2 cells or treating them with Fe^2+^ (ferrous sulfate) alone did not cause cell death as measured by LDH release (**Figure 5F**). Albumin treatment increased cell death comparably in scramble siRNA and FtH1 knocked down cells (**Figure 5F**). However, concomitant treatment with Fe^2+^ and albumin resulted in most cell death in the scramble siRNA treated cells and was significantly more in the FtH1 knocked down cells (**Figure 5F**). Collectively these data indicate that loss of FtH1 exacerbates proximal tubular epithelial cells injury and death in glomerulonephritis.

**Figure 6.**
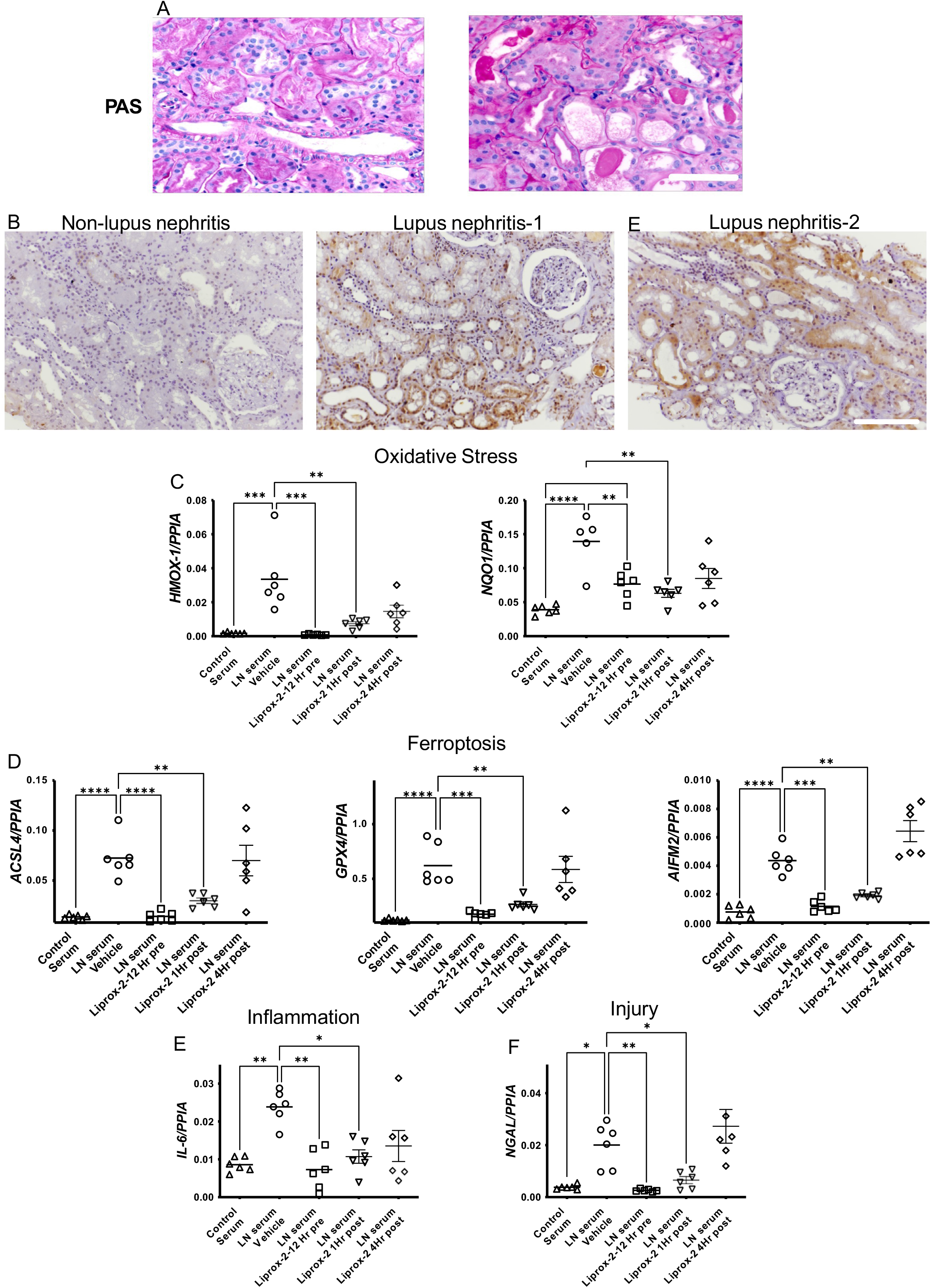
Ferroptosis is observed in human lupus nephritis and lupus nephritis patients’ serum-induced ferroptosis in proximal renal tubules is mitigated by Liproxstatin-2. Compared to non-lupus nephritis controls, PAS-stained human biopsies from patients with class IV lupus nephritis show acute tubular injury with tubular dilatation, epithelial attenuation, and casts (A). Lupus nephritis kidneys stain more intensely for 4HNE, lipid peroxidation and the ferroptosis marker (B). Most of the staining is in the tubular segments, the area of iron accumulation. Representative images from two individual patient kidneys with or without lupus nephritis are shown. Scale bar = 50 μm, 100 μm. HK2 cells, (a human proximal tubular cell line) were treated with vehicle (DMSO) or 100 nM Lipoxstatin for 12 hrs and then cultured with 5% normal donor serum or serum from patients with class IV lupus nephritis. In some experiments Liproxstatin-2 was added 1 or 4 hrs after addition of the serum. All studies were terminated 24 hrs after serum addition. Compared to normal serum, lupus nephritis serum induced a significant increase in gene expression of HMOX-1, NQO1 (C) (Oxidative Stress) and ACSL4, GPX4 and AIFM2 (D) (Ferroptosis), IL-6 (Inflammation) (E) and NGAL (injury) (F). All these pathological parameters were significantly reduced by Liproxstatin-2 pretreatment or by adding it 1 hr after serum exposure. Lirpoxstatin-2 did not have any benefit when added 4 hrs after exposure to lupus nephritis serum. Data was analyzed using 2-way ANOVA with Tukey’s multiple comparison test. Only significant values are showed for comparison. Each dot represents an individual donor. Data is presented as mean. *P < 0.05, **P < 0.001, *** < 0.0001.

### Tubular ferroptosis is a prominent feature in human lupus nephritis and can be reversed *in vitro* by Liproxstatin-2, a new generation ferroptosis inhibitor

After confirming the occurrence of ferroptosis in murine models of lupus nephritis, we next determined whether ferroptosis is also a feature of human lupus nephritis. Compared to non-nephritic controls, PAS-stained kidney biopsies of Class IV lupus nephritis patients showed regions of acute tubular injury with dilated tubules and casts **(Figure 6A)**. Non-nephritic control and lupus nephritis (Class IV) kidney biopsies were stained for the expression of 4HNE and ACSL4. We observed a higher expression of 4HNE (lipid peroxidation, ferroptosis) in the tubular segments of human class IV lupus nephritis biopsies **(Figure 6B)**. This was similar to the 4HNE staining pattern observed in nephritic mice (**Figure 3G-H)**. Similarly, ACSL4 (a marker of ferroptosis) staining was primarily observed in the tubular segments of the kidney biopsies (**Supplemental Figure 7**). These data indicate the occurrence of ferroptosis in the kidney tubules of human Class IV lupus nephritis. Since ferroptosis was primarily observed in the tubular segments, we evaluated the effect of lupus nephritis patient’s serum on HK-2 cells, a proximal tubular cell line. Compared to healthy controls, incubation of HK-2 cells with 5% lupus nephritis patients’ serum (Class IV) significantly increased markers of oxidative stress (**Figure 6C**) ferroptosis (**Figure 6D**). Lupus nephritis serum also significantly increased inflammation and injury in HK-2 cells (**Figure 6E-F**). All these pathological features induced by lupus nephritis serum were significantly mitigated by Liproxstatin-2 (**Figure 6C-F)**. Importantly, Liproxstatin-2 demonstrated therapeutic benefit when administered an hour after the addition of lupus nephritis serum (**Figure 6C-F)**. Liproxstatin-2 did not offer any benefit when administered 4 hrs after the addition of lupus nephritis serum (**Figure 6C-F)**.

## Discussion

The presence of tubular immune complexes and proximal tubular epithelial cell injury in nephritic mice supports the ongoing notion that tubular injury plays a critical role in the pathogenesis of lupus nephritis. Our study provides evidence of ferroptosis in the tubular compartment of human and murine lupus nephritis. Furthermore, using the *FtH*^*PT−/−*^ mice and cells, we link impaired iron sequestration to increased tubular injury in primary glomerulonephritis. Finally, we demonstrate that lupus nephritis serum components directly induce ferroptosis in human proximal tubular epithelial cells, which can be reversed by a new generation ferroptosis inhibitor, Liproxstatin-2. Thus, our study identifies tubular iron accumulation and ferroptosis as a possible link between glomerular injury and ensuing tubulointerstitial inflammation in lupus nephritis.

The expression of ACSL4, a key ferroptosis executioner, was increased in nephritic human biopsies and kidneys of diseased and nephritic mice. Functionally, ACSL4 ligates arachidonic acid with CoA to generate arachidonoyl-CoA, which is subsequently conjugated to phosphatidylethanolamine mediated by lysophosphatidylcholine acyltransferase (LPCAT)(20). These esterified phosphatidylethanolamine conjugates are oxidized by lipoxygenases to generate lipid hydroperoxides, the proximate executors of ferroptosis(39, 42). Peroxidized phospholipids are counteracted by GPX4 in the presence of glutathione, a cofactor of GPX4 (43), that reduces toxic lipid peroxides to nontoxic lipid alcohols thereby inhibiting ferroptosis(24). Diseased, female, and male kidneys of two models of lupus nephritis have decreased expression of GPX4, a new observation in this disease. We also report that *Gclc* and *Gclm*, the gene products that make up the enzyme GCL, are significantly reduced in the diseased kidneys of both lupus-prone female and male mice, indicating impaired GCL activity. Reduced GCL activity directly inhibits GSH synthesis(44) and may account for the observed reduction in GPX4. Our observations may be of direct relevance in understanding the pathophysiology of lupus nephritis since lupus patients have decreased levels of GCL and GSH(45).

Our lipidomics data further validate the sensitization of renal tissue towards ferroptosis in lupus nephritis. We observed increased PE(P-18:0/20:4), PE(P-18:1/20:4), and PE(P-18:2/20:4) in the kidneys of nephritic mice. Phosphatidylethanolamines are the major storage depots for arachidonic acid in cells(46). They can be hydrolyzed to liberate arachidonic acid (47), the substrate for ACSL4. In support, we found a significant increase in esterified sn-2 chain of PE (P-18:0/22:4). Suppression of esterification of PE by genetic or pharmacological inhibition of ACSL4 acts as a specific anti-ferroptotic rescue pathway(39). Thus, our observations identify ACSL4 as a potential target for intervention in lupus nephritis. Relevant to this, only one study has described intra-renal lipid profiles in lupus nephritis(48) and proposed them as a potential early biomarker for lupus nephritis.

To highlight the importance of excess iron accumulation in tubular pathology secondary to glomerular injury, we utilized transgenic mice that are deficient for FtH1 in their proximal tubules. FtH1 deficiency in proximal kidney tubules (*FtH*^*PT−/−*^) worsens kidney pathology following rhabdomyolysis and unilateral ureteric obstruction(28, 49), but the influence of FtH1 on the outcomes of tubular injury following glomerulonephritis has not, to our knowledge, been reported previously. Mechanistically, FtH1 oxidizes labile or bioactive ferrous (Fe^2+^) to neutral ferric (Fe^3+^) form and sequesters it safely within its core(33, 34), thereby limiting iron-induced ROS. We demonstrate that FTH1 deficiency increases tubular injury, secondary to glomerulonephritis, and is independent of the extent of glomerular injury. Together with our in vitro findings that, concomitant treatment with Fe^2+^ and albumin exacerbates cell death in FtH1 knocked down PTECs cells (**Figure 7F**) our proof-of-concept experiments identify dysregulation of tubular iron metabolism as a susceptibility factor in glomerulonephritis and support the human observations that following glomerulonephritis, the extent of tubular injury is independent of glomerular pathology(9, 50). In addition, they identify FtH1 inducing compounds as a potential adjunct therapy for lupus nephritis(19, 34).

Finally, we demonstrate that serum components of class IV lupus nephritis patients can directly induce ferroptosis in human proximal tubular cells, a new observation in the field. Remarkably, the prophylactic and therapeutic treatment with Liproxstain-2, a novel 2^nd^ generation ferroptosis inhibitor significantly reduced lupus nephritis patients’ serum-induced ferroptosis and injury to levels equivalent to or lower than non-lupus nephritis controls. We choose this population as it involves diffuse lupus nephritis involving 50% or more of glomeruli, which indicates severe LN requiring more-intensive therapy. A recent study identified serum autoantibody from SLE patients and type I interferon independently induced neutrophil ferroptosis(27). This was inhibited by liproxstatin-1 and the iron chelator deferoxamine. While our study does not identify whether autoantibodies of different specificities or cytokine milieu are needed for proximal tubular epithelial cells ferroptosis, it provides a new approach to inhibit proximal tubular epithelial cells pathology in lupus nephritis.

### Limitations and future directions

The observations made here with a limited number of human samples are supported by two murine models of spontaneous lupus nephritis with different etiologies. The lack of in vivo data using Liproxstatin-2 is a limitation of our study, but these novel observations warrant future investigations in long term spontaneous animal models of lupus nephritis. Along with the work of Li et. al (27) that demonstrate the efficacy of ferroptosis inhibition in SLE, such future studies will reinforce the utility of ferroptosis inhibitors as a powerful adjunct therapeutics and aid current therapies to increase rates of remission in this devastating disease. As discussed above Ferroptosis plays an essential role in multiple forms of kidney injury and diseases, but the mechanistic characterization of tubular cell pathology in lupus nephritis is lagging. Our study identifies a novel druggable pathological mechanism contributing to tubular pathology during the evolution of lupus nephritis.

## Author Contribution

YS conceptualized the work and wrote the original draft. AAA, DD, AE, and YS did most of the experiments. AA, LAS, and NDD performed lipidomics and mass spectrometric studies. WC and CA scored the kidney pathology. LM generated the (NZW X BXSB) F1 mice, and BM provided input for iron biology. BP and MC provided Liproxstatin-2 and helped design the in-vitro studies. MS helped with the patient studies. SB helped generate the *FtH*^*PT−/−*^ mice. All authors helped edit and approve the final version of the manuscript.

## Acknowledgments

The authors are thankful to Dr. Anupam Agarwal (University of Alabama) for his input in designing the experiments in *FtH*^*PT−/−*^ mice and Dr. Volker Haase (Vanderbilt University) for providing the PepckCre mice.

## Supplemental Methods for

## Supplemental Methods

### Immunohistochemistry

Paraffin-embedded human biopsies or mouse kidney sections (3 μm) were stained for acyl-CoA synthetase long-chain family member 4 (ACSL4) or 4-hydroxynonenal (4-HNE) using standard protocols. Sections were deparaffinized in xylenes, rehydrated in a series of ethanol rinses (100%–70%), and washed in distilled water. Sections were then incubated in 3% H_2_O_2_ in methanol for 20 minutes. After antigen retrieval by heating in citrate buffer and treating with avidin and biotin for 15 minutes each (Avidin/Biotin Blocking Kit; Vector Laboratories), the sections were incubated in blocking buffer containing 10% donkey serum in 0.1 M sodium phosphate buffer pH 7.4 (phosphate buffer) at room temperature for 30 minutes. Next, sections were incubated with anti-ACSL4 (1: 500, in 1% BSA/phosphate buffer; sc-271800, Santa Cruz) or 4-HNE antibody (1:1000 in 1% BSA/phosphate buffer; Abcam, Boston, MA) overnight at 4°C. After three washes with phosphate buffer for 5 minutes each, sections were incubated with biotinylated donkey anti-mouse or donkey anti-goat secondary antibody (1:400 dilution; Vector Laboratories) for 1 hour. After three washes in phosphate buffer, sections were incubated with ABC ready-to-use reagent (VECTASTAIN Elite ABC Kit; Vector Laboratories) for 30 minutes. After another three washes with phosphate buffer, the sections were incubated with 3,3′-diaminobenzidine for 5 minutes, followed by washing with distilled water. Finally, the sections were counterstained with 1% methylene blue solution, washed in water, and dehydrated with xylene. Sections were imaged using a Nikon microscope.

### Immunofluorescence and detection of immune complex

Three to five micron cryostat-cut kidney sections were used for the immunofluorescence detection of α smooth muscle actin (αSMA), F4/80+ macrophages, CD4+ T cells and immune complexes. Detailed method is provided in the supplemental methods sections. Briefly, the kidney sections were air-dried and incubated in PBS with 0.3% TritonX-100/10% horse serum. After washing with PBS, FC receptors were blocked with anti-CD16/32 (2.4g2, ThermoFisher, Waltham, MA USA) antibody. This was followed by incubation with FITC-labeled anti-αSMA (1A4, 1:30; Sigma, St. Louis, MO), PE-labeled F4/80 (BM8, 1:20; ThermoFisher), PE-labeled CD4 (GK1.5, 1:20; ThermoFisher) in 10% horse serum/PBS for 1.5 hrs. The sections were then washed in PBS and mounted with ProLong Gold antifade agent with DAPI (ThermoFisher).

To examine glomerular immune complex deposits, 5 mm snap-frozen kidney sections were fixed in acetone, washed in PBS, incubated in 3 % hydrogen peroxide for 10 min, and rinsed in PBS. Slides were incubated in FITC–conjugated monoclonal anti-IgG antibody (Santa Cruz, Dallas, Tx) at a 1:50 dilution in PBS for 1 hr at room temperature. Slides were then rinsed in PBS and mounted with a coverslip using Vectashield hard set medium (Vector H-1400, Vector Laboratories, Burlingame, CA). Fluorescence was detected using a Nikon microscope.

### Western Blotting

Snap-frozen tissue sections were homogenized in Tris-Triton tissue lysis buffer containing complete protease inhibitor cocktail (Halt Protease and Phosphatase Inhibitor Cocktail; Thermo Fisher Scientific, Rockford, IL) using a Dounce Homogenizer. GPX4 and ACSL4 expression were measured in whole kidney lysates. Protein content in the homogenate was estimated using the Pierce BCA protein estimation kit (Thermo Fisher Scientific). Twenty micrograms of protein per sample were loaded on a 4-12% NuPage Bis-Tris gel under reducing conditions. The resolved proteins were transferred onto a nitrocellulose membrane (LI-COR Biotechnology, Lincoln, NE) and probed with rabbit monoclonal anti-GPX4 (1:1000, ab125066, Abcam), mouse monoclonal anti-ACSL4 (1:100, sc-271800, Santa Cruz), rabbit monoclonal anti-FtH1 (1:1000, ab183781, Abcam) and mouse monoclonal anti-βactin (1:7000, ab8226, Abcam) antibodies. The primary antibodies were detected using donkey anti-rabbit Alexa 800 and donkey anti-mouse Alexa 800 antibodies (LI-COR). Mouse monoclonal anti-β-actin (Abcam) was used as the loading control and detected using donkey anti-mouse Alexa 680 antibody (LI-COR). Data were quantified using densitometry software (LI-COR).

### RT-PCR

For RNA isolation, frozen tissues or live cells were resuspended in RLT buffer (Qiagen, Valencia, CA) and homogenized using the TissueLyser system or Qiashredder (Qiagen). RT was performed as described in our previous publication(1). Total RNA from homogenates was purified using the RNAeasy mini kit (Qiagen) following the manufacturer’s instructions. Then, 0.2-1 μg of RNA was used to synthesize cDNA using the iScript cDNA synthesis kit (Bio-Rad, Hercules, CA). The cDNA template was mixed with iTAQ SYBR green universal supermix (Bio-Rad), and quantitative PCR was performed on a CFX Connect system (Bio-Rad). Human or mouse predesigned gene primers were purchased from Bio-Rad. Human or mouse *PPIA* was amplified in parallel and used as the reference gene in quantification. Data are expressed as relative gene expression and were calculated using the 2^[−ΔC(T)] method.

### Cell Culture and siRNA knockdown studies

For gene knockdown studies, silencer select siRNA for human FtH1 and negative control siRNA (Catalog number: 4392421 and 4390843 respectively, ThermoFisher Scientific) were used. HK-2 cells (2 × 10^5^/well) grown in the recommended medium were transfected with the FtH1 and negative control siRNA, respectively, using Opti-MEM reduced serum medium and Lipofectamine RNAiMAX Transfection Reagent (ThermoFisher Scientific) in 12 well plates as per the manufacturers instructions. Gene knockdown efficiency was measured after 52 hrs by RT-PCR. At this time point medium was changed and cells were incubated for 12 hrs with vehicle, 200 μM ferrous sulfate, 20 mg/mL human albumin or a mixture of 200 μM ferrous sulfate and 20 mg/mL human albumin. Lactate dehydrogenase (LDH) content in the cell culture supernatant was used to measure cell death (Cayman Chemical).

### Preparation of membrane fractions and liquid chromatography-mass spectrometric analysis of renal lipids

For each animal, 10 mg kidney tissue was washed with PBS before being subject to homogenization with an Omni TH homogenizer (Warrenton, VA) in tissue protein extraction reagent (TPER) (Thermo Fisher Scientific; Waltham, Massachusetts) supplemented with Halt protease and phosphatase inhibitors (Thermo Fisher Scientific). The lysate was incubated on ice for 30 minutes while vortexing every 10 minutes. The lysates were then subject to centrifugation at 13,000 rpm for 10 minutes using a tabletop centrifuge (Thermo IEC) before the supernatant was subject to ultracentrifugation for 30 minutes at 34,000 rpm at 4º C using an SW55.1 rotor and Optima L-90K ultracentrifuge (Beckman Coulter; Schaumburg, IL). The pellet was reconstituted in TPER and then sonicated twice for 5-second intervals. A BCA assay was performed to determine protein concentration.

Lipids were extracted with a modified Bligh-Dyer protocol(2). Two milliliters of methanol, 800 uL of dichloromethane, 20 uL of lipid membranes, 980 ul of water, and 50 ul of 1:5 diluted EquiSplash Lipidomix heavy isotope Standards (consisting of 20 ug/L each of a mixture of 13 deuterated lipids, Avanti Polar Lipids, Inc., AL, USA) were combined in 13×100 mm glass screw-cap tubes. The mixture was left at room temperature for thirty minutes. 1 mL water and 0.9 mL methylene chloride were added and tubes were gently inverted 10 times and centrifuged at 200 g for 10 minutes. After centrifugation, the lower layer was collected. The samples were extracted with 800 ul of dichloromethane, vortexed, centrifuged before collecting the lower layer. Finally, the organic phase was dried under nitrogen gas and reconstituted in 50 uL of methanol. Targeted liquid chromatography-mass spectrometric analysis was performed on a QTRAP 6500+ triple quadrupole mass spectrometer (AB SCIEX) and a Nexera liquid chromatography system (Shimadzu). The column used was a Phenomenex luna 3μm 2.1×100 mm column with mobile phase A 1 mM ammonium acetate in 7/93 dichloromethane/acetonitrile and mobile phase B consisting of 1 mM ammonium acetate in 50/50 water/acetonitrile pH 8.2. The gradient was 17.5 minutes long with a 5 ul injection volume. About 1,200 lipids were detected. Differentially regulated lipids of different classes were compared between non-nephritic and nephritic kidneys, and p < 0.05 was considered significant. A heatmap was used to visualize the results.

---

**Supplemental Figure 1.**
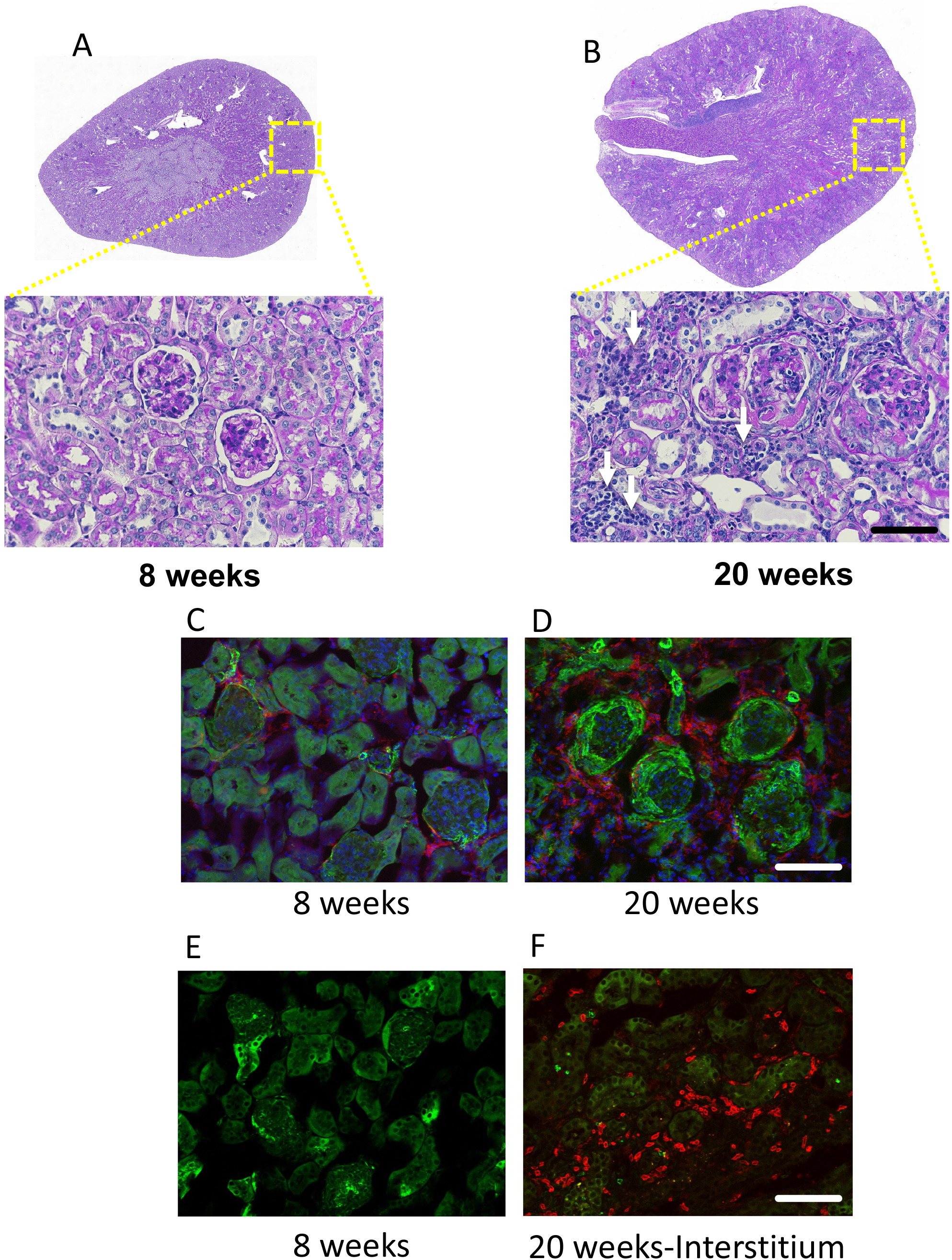
Tubulointerstitial injury in lupus nephritis is accompanied by immune cell infiltration composed of macrophages and CD4 T cells. The kidneys of 8- and 20-week-old female MRL/lpr mice were analyzed by morphometry and immunofluorescence. Representative morphology by Periodic acid-Schiff (PAS)) of kidneys showed severe changes in glomeruli and tubulointerstitium with disease progression: endocapillary proliferation, infiltration of immune cells, formation of wire loops and crescents, atrophy and sclerosis is evident with age (Fig 1**A-B** and inset). White arrows point to tubulointerstitial infiltrates. At 20-weeks of age alpha smooth muscle actin positive cells are seen surrounding the glomeruli, with F4/80 positive macrophages in the periglomerular and peri-tubular regions(Fig 1**C-D**). Similarly, CD4+ T cells are seen in abundance in the periglomerular and peri-tubular regions of the nephritic kidneys (Fig 1**E-F**) Collectively this indicates that lupus nephritis is associated with severe glomerular and tubulointerstitial injury.

**Supplemental Figure 2.**
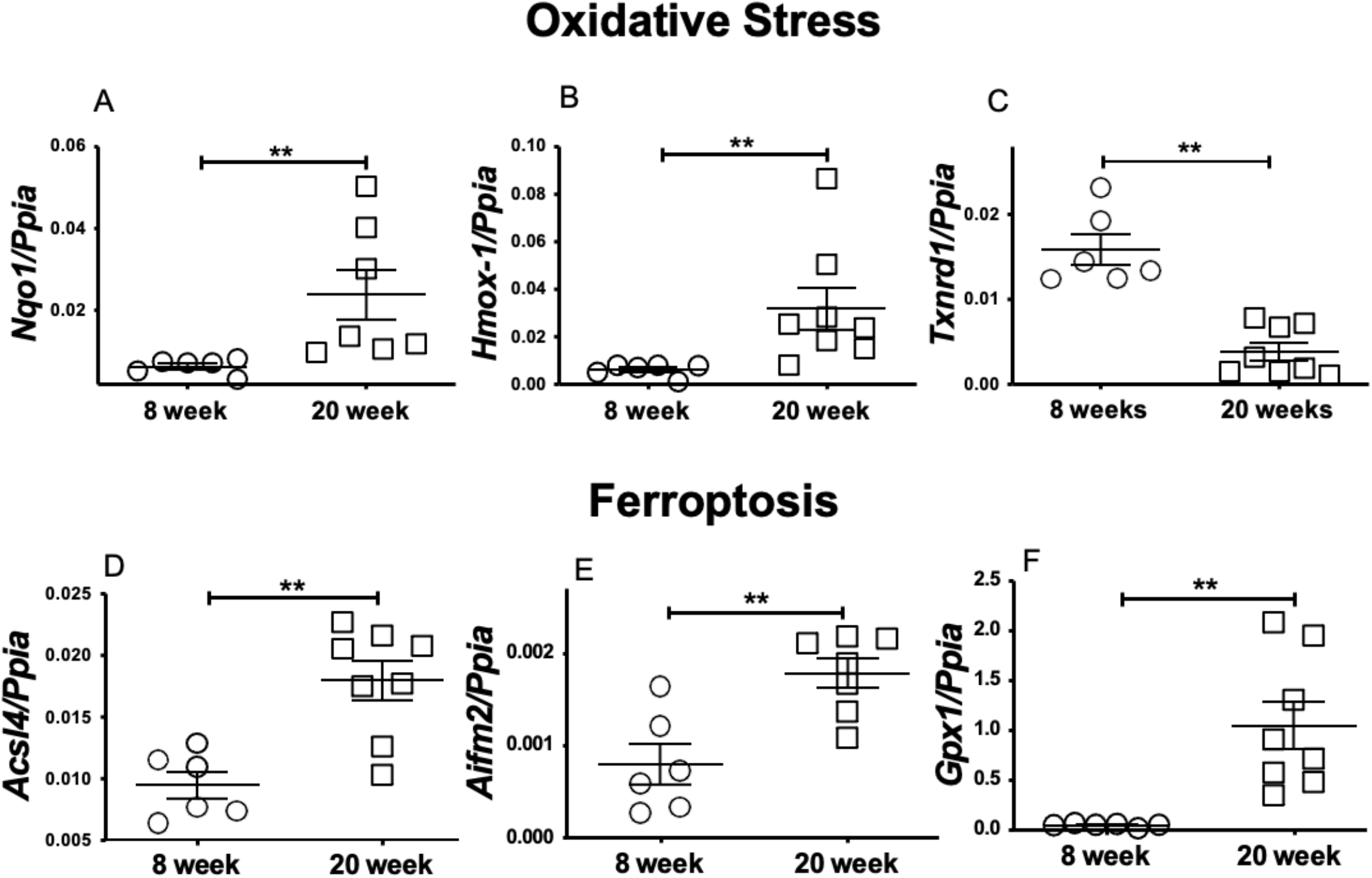
Lupus nephritis kidneys display oxidative stress and a ferroptosis gene signature. Compared to 8-week-old MRL/lpr females, 20-week-old nephritic females have increased oxidative stress as indicated by significantly higher gene expression of *Nqo1* **(A)**, *Hmox-1* **(B)** and lower expression of *Txnrd1* **(C)**. Increased iron accumulation (Figure 2) and oxidative stress in nephritic mice was associated with an increase in renal gene expression of ferroptosis markers like *Acsl4* **(D)**, *Aifm2* **(E)**, *Gpx1* **(F)**. Statistical significance was determined by 2-tailed Mann-Whitney test. Data are plotted as mean ± SEM. **P < 0.001.

**Supplemental Figure 3.**
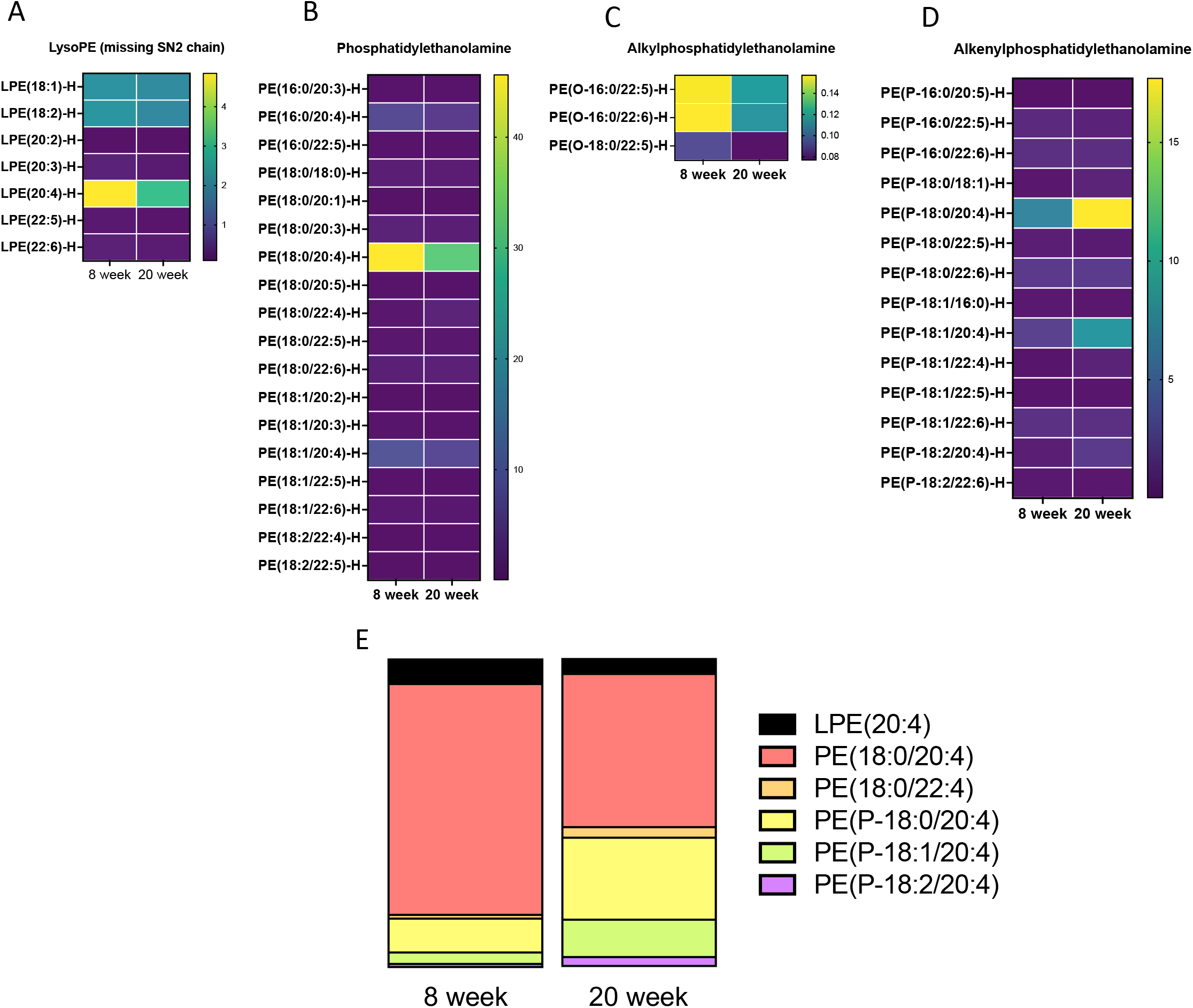
Differential lipid profile in the kidneys of non nephritic (8-week-old) and nephritic (20-week-old) mice. 70-80 mg kidneys from 8-week-old (non nephritic) and 20-week-old (nephritic) MRL/lpr females were used to isolate lipids from the plasma membrane. Subsequently, the isolated lipids were subjected to semi-targeted liquid chromatography-mass spectrometry (LC-MS) to identify and quantify their lipid profile and content. Heat maps were generated by normalizing the concentration of individual lipids within each class and category to total lipid concentration within each kidney (n=6 each). Heat maps of significantly different lysophosphotidylethanolamine: LPE, phosphotidylethanolamine: PE, alkyl-phosphatidylethanolamine: PE(O) and alkenylphosphatidylethanolamine: PE(P) classes of lipids show a clear differential expression in individual lipids within and between different classes **(A-D)**. Intrarenal fingerprint obtained by averaging most differentially expressed lipids in LPE, PE, and PE(P) in non nephritic and nephritic kidneys **(E)**.

**Supplemental Figure 4.**
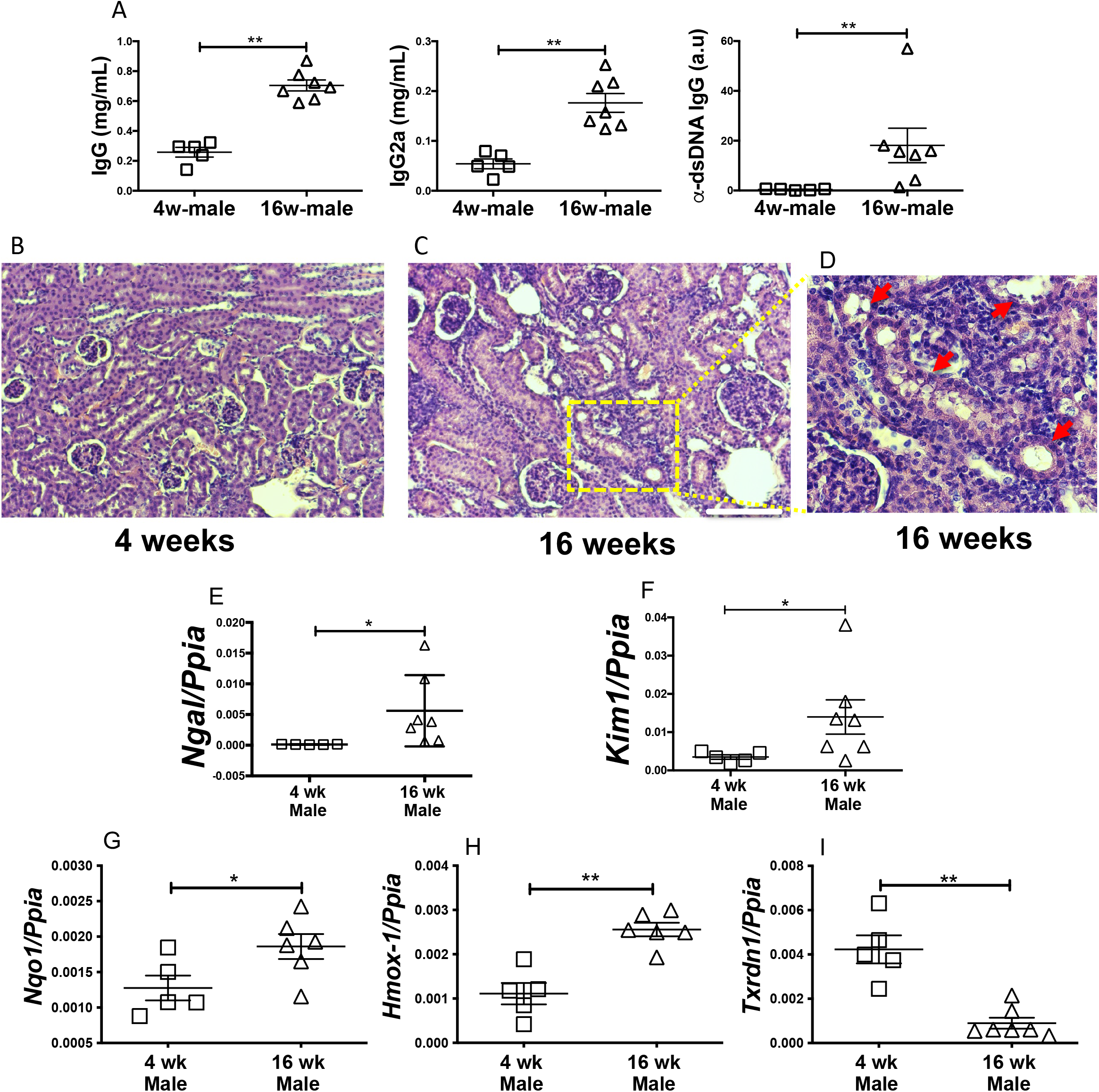
Tubular injury and oxidative stress signature are a feature in a male mouse model of lupus nephritis with different etiology. The (NZW X BXSB) F1 male mice carry two copies of TLR7 and develop progressive glomerulonephritis starting from 14 weeks of age. At 16 weeks of age, they have high titers of IgG and anti dsDNA antibodies (**A**). Compared to 4-week-old, the renal histology (H&E) of 16-week-old males shows glomerular hypertrophy, infiltration as well as injured renal tubules with dilatation, luminal debris (red arrows) (**B-D**). Scale bar = 50 μm. Representative images are shown. Proximal tubular injury markers *Ngal* (**E**) and *Kim1* (**F**), were significantly increased in 16-week-old mice. Thus, a TLR7 driven, male model of lupus also develops glomerular and tubular injury. Injury to the kidneys is associated with increased oxidative stress as indicated by an increase in *Nqo1* (**G**), *Hmox1* (**H**) and a decrease in *Txrdn1* (**I**). Statistical significance was determined by 2-tailed Mann-Whitney test. Data is presented as mean ± SEM.*P < 0.05, **P < 0.001.)

**Supplemental Figure 5.**
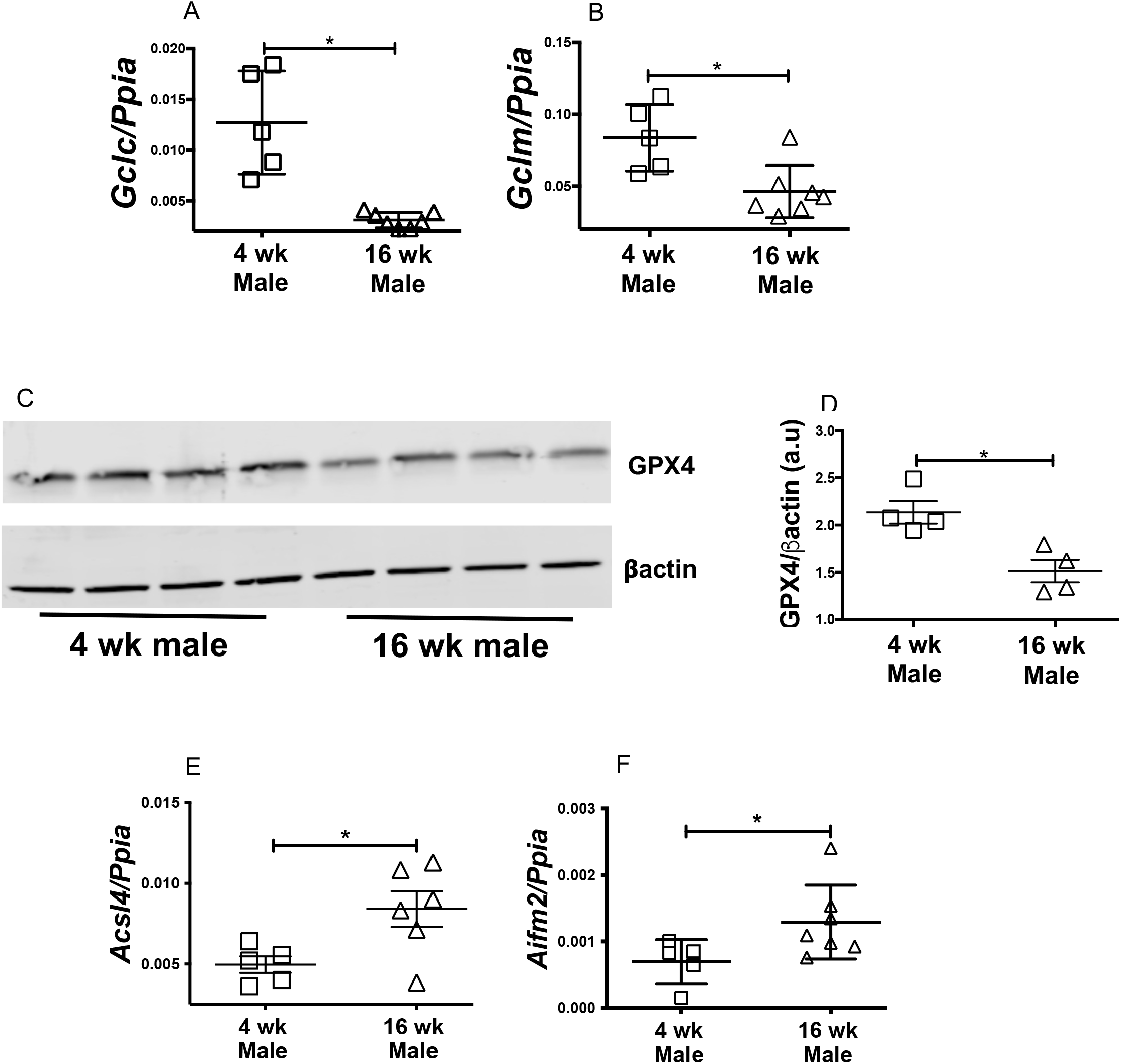
Impaired glutathione biosynthesis program and ferroptosis signature in (NZW X BXSB) F1 male mice. The (NZW X BXSB) F1 male carry two copies of TLR7 and develop progressive glomerulonephritis starting from 14 weeks of age. 16-week-old nephritic males show significantly lower expression of the catalytic subunit *Gclc* (**A**) and modifier subunit *Gclm* (**B**), of glutamate cysteine ligase (GCL), the activity of which determines de novo glutathione synthesis. Compared to 4-week-old males, 16-week-old nephritic males had significantly lower expression of GPX4, the glutathione dependent ferroptosis inhibitor (**C and D**). Additionally, other markers of ferroptosis like *Acsl4* (**E**) and *Aifm2* (**F**) were also significantly elevated in 16-week-old nephritic males. Statistical significance was determined by 2-tailed Mann-Whitney test. Data are plotted as mean ± SEM. *P < 0.05. Collectively these data indicate that ferroptosis is a feature in lupus prone mice driven by different etiologies.

**Supplemental Figure 6.**
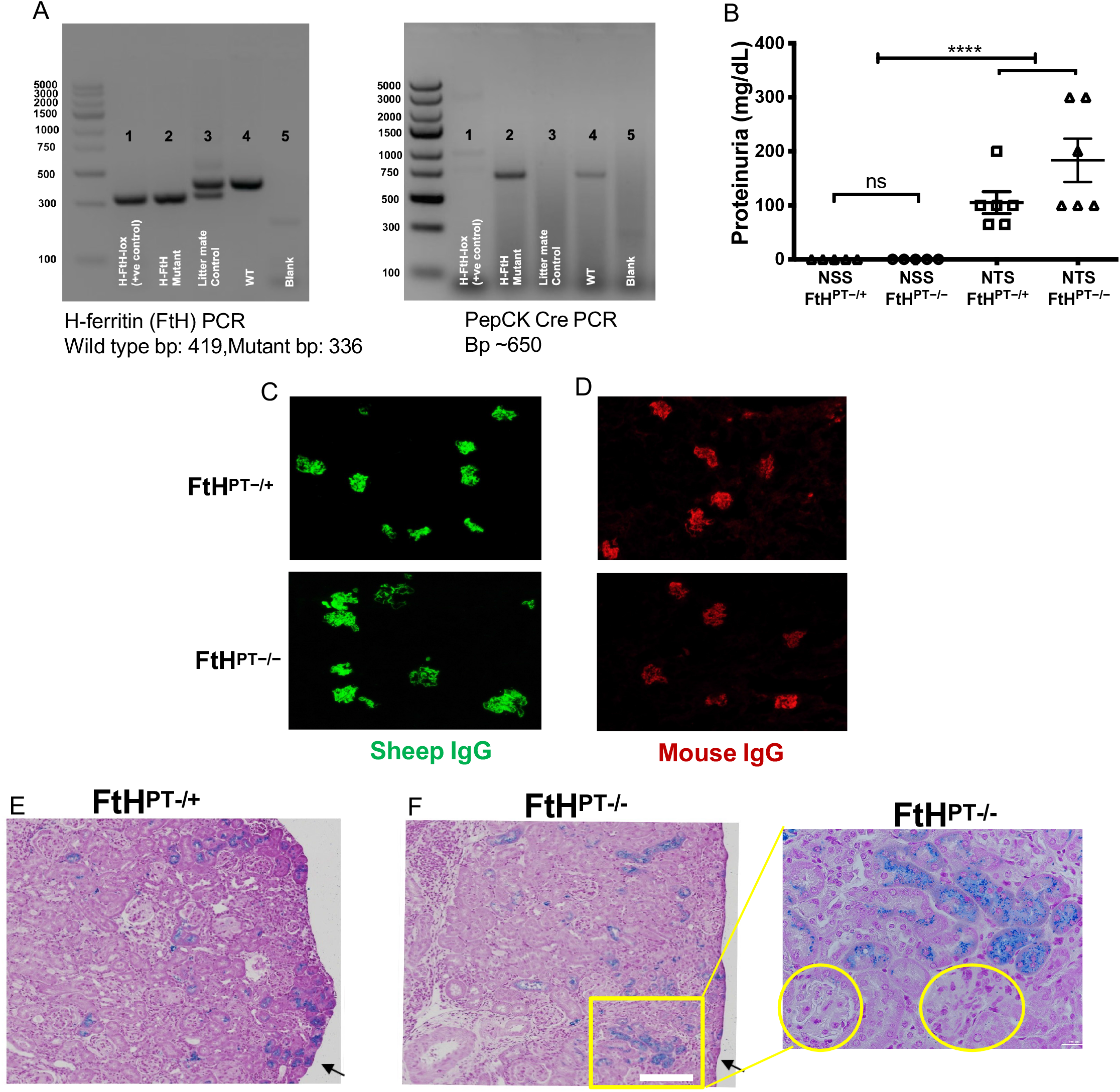
Comparable proteinuria, glomerular immune complexes and tubular iron deposits in wild type mice and mice selectively deficient for FtH1 in their proximal renal tubules. Genotype of *FtH*^*PT−/−*^ and *FtH*^*PT−/+*^ mice used in this study (**A**). Lanes 1) FtH flox/flox (H-FtH-lox)→ positive control for loxed allele, 2) *FtH*^*PT−/−*^ (H-FtH mutant) → Mouse deficient for FtH1 in proximal renal tubules, 3) *FtH*^*PT−/+*^ → Litter mate control, 4) WT (Wild type) 5) Blank. 12-week-old female FtH^PT-/-^ or FtH^PT-/+^ (littermate controls) mice, were pre-sensitized with 100 ug sheep IgG in CFA, and four days later were injected i.v., with 100 uL **normal sheep serum (NSS) or nephrotoxic sheep serum (NTS)**. Kidneys were analyzed 14 days later. NSS did not cause any glomerulonephritis in FtH^PT-/-^ or FtH^PT-/+^ mice (**B**). In contract NTS caused comparable glomerular injury in both FtH^PT-/-^ and FtH^PT-/+^ mice as measured by proteinuria (**B**). We also observed comparable anti-sheep (heterologous) and anti-mouse (autologous) glomerular immune complex deposits (**C**) and (**D**). Similar Perls detectable iron deposits were detected in the tubular segments of FtH^PT-/+^ and FtH^PT-/-^ mice, whereas glomeruli (yellow circles) were devoid of observable iron deposit (**E-F and inset**). Scale bar 100 uM and 50 uM. Data was analyzed using 2-way ANOVA with pos-hoc analysis and represented as mean ± SEM. ***P < 0.0001.

**Supplemental Figure 7.**
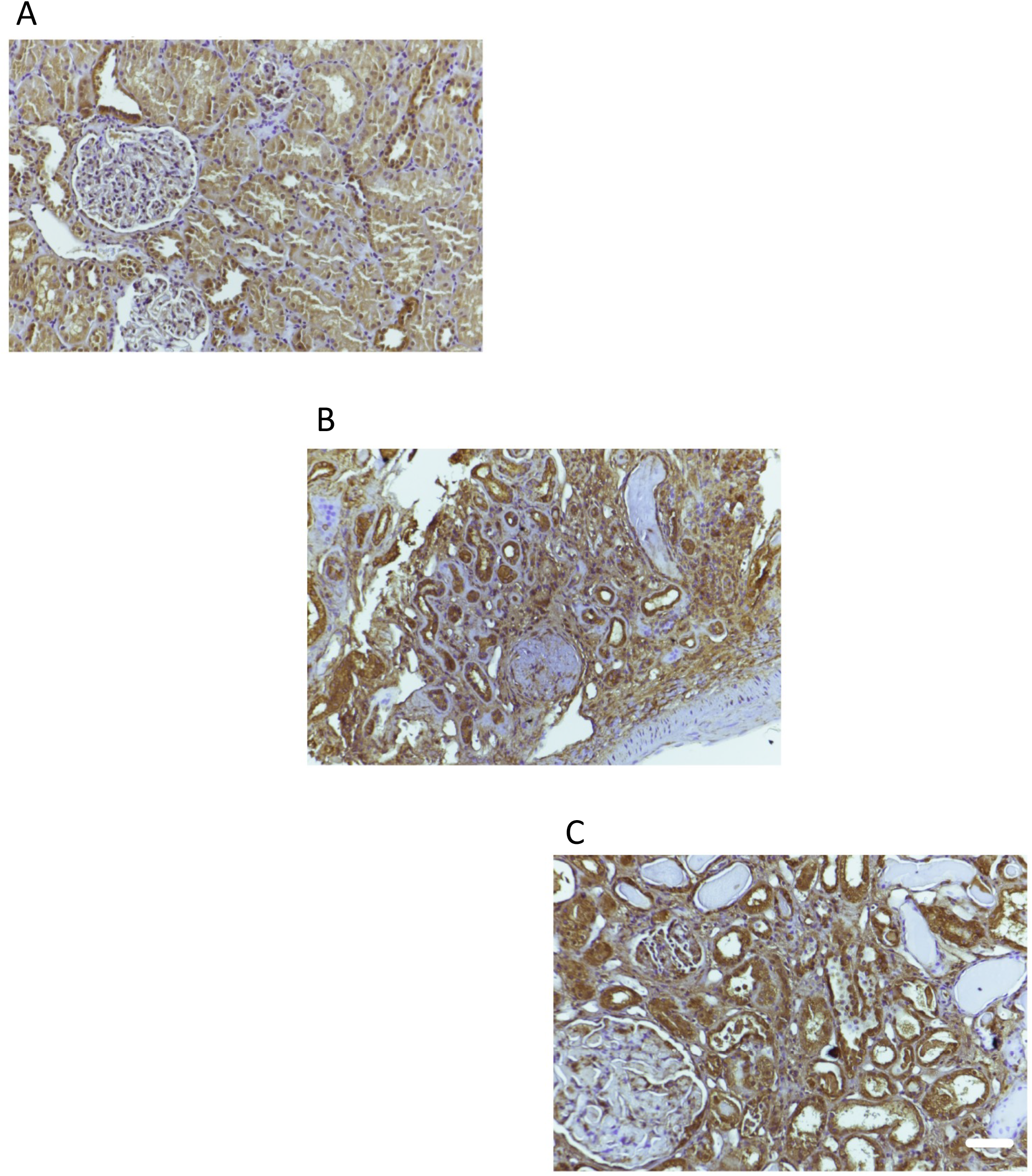
Ferroptosis marker ACSL4 staining in human non-lupus and lupus nephritis kidneys. Non lupus patient’s kidney (**A**) and 2 individual lupus nephritis kidneys with class IV lupus nephritis were stained for ACSL4. ACSL4 expression was observed in some of the renal tubules of non-lupus patient stained (**A**). However, most of the renal tubules of patients with class IV lupus nephritis were positive and showed more intense staining (**B-C**). Scale bar = 50 μm. The glomeruli in all the samples did not show show any appreciable staining. Increased ACSL4 expression (ferroptosis promoter) aligned well with 4-HNE staining observed in lupus nephritis kidneys (Figure 8)

